# GeneCOCOA: Detecting context-specific functions of individual genes using co-expression data

**DOI:** 10.1101/2024.06.27.600936

**Authors:** Simonida Zehr, Sebastian Wolf, Thomas Oellerich, Matthias S. Leisegang, Ralf P. Brandes, Marcel H. Schulz, Timothy Warwick

## Abstract

Extraction of meaningful biological insight from gene expression profiling often focuses on the identification of statistically enriched terms or pathways. These methods typically use gene sets as input data, and subsequently return overrepresented terms along with associated statistics describing their enrichment. This approach does not cater to analyses focused on a single gene-of-interest, particularly when the gene lacks prior functional characterization. To address this, we formulated *GeneCOCOA*, a method which utilizes context-specific gene co-expression and curated functional gene sets, but focuses on a user-supplied gene-of-interest. The co-expression between the gene-of-interest and subsets of genes from functional groups (e.g. pathways, GO terms) is derived using linear regression, and resulting root-mean-square error values are compared against background values obtained from randomly selected genes. The resulting *p* values provide a statistical ranking of functional gene sets from any collection, along with their associated terms, based on their co-expression with the gene of interest in a manner specific to the context and experiment. *GeneCOCOA* thereby provides biological insight into both gene function, and putative regulatory mechanisms by which the expression of the gene-of-interest is controlled. Despite its relative simplicity, *GeneCOCOA* outperforms similar methods in the accurate recall of known gene-disease associations. *GeneCOCOA* is formulated as an R package for ease-of-use, available at https://github.com/si-ze/geneCOCOA.

**Author summary:** Understanding the biological functions of different genes and their respective products is a key element of modern biological research. While one can examine the relative abundance of a gene product in transcriptomics data, this alone does not provide any clue to the biological relevance of the gene. Using a type of analysis called co-expression, it is possible to identify other genes which have similar patterns of regulation to a gene-of-interest, but again, this cannot tell you what a gene does. Genes whose function has previously been studied are often assembled into groups (e.g. pathways, ontologies), which can be used to annotate gene sets of interest. However, if a gene has not yet been characterized, it will not appear in these gene set enrichment analyses. Here, we propose a new method - *GeneCOCOA* - which uses co-expression of a single gene with genes in functional groups to identify which functional group a gene is most similar too, resulting in a putative function for the gene, even if it has not been studied before. We tested *GeneCOCOA* by using it to find gene-disease links which have already been scientifically studied, and showed that *GeneCOCOA* can do this more effectively than other available methods.

## Introduction

Advances in sequencing technology have decreased the costs and increased the accuracy of transcriptome profiling [1]. This has resulted in an abundance of datasets generated from a wide variety of experimental conditions, many of which are made publicly available [2–4]. As such, interrogation of public sequencing data has become an increasingly important step in research focused on a specific gene or gene product of interest. Normally, this is limited to detecting whether the gene-of-interest is expressed in a given dataset or whether the expression of the gene changes in a particular experimental condition [5]. However, this approach does not supply insight into any potential functions of the gene-of-interest in the data, or any regulatory mechanisms which might govern expression of the gene.

Functional enrichment analyses carried out in the course of differential gene expression analysis usually relies upon the input of one or more gene sets which are derived throughout the course of the analysis (e.g. differentially expressed genes) [6]. Curated associations between each gene and sets of annotations such as ontologies [7], pathways [8,9] and diseases [10] are then computed. These associations are subsequently statistically analyzed for overrepresented terms, considering the size of the input gene set, the number of genes associated with the given term, and enrichment in hits compared to an appropriate background gene set [11–15]. The outcome of these analyses is a list of terms stratified by statistical values such as *p* value, adjusted *p* value, precision and recall. Results from these approaches have the potential to inform future research directions and wet-lab experiments. However, they cannot provide insight into the functional relevance of individual genes, especially when genes lack prior functional characterization.

One approach that can be used to examine potential function of an individual gene-of-interest (GOI) is to model the expression of the GOI against the expression of other genes present in a given dataset, in a co-expression analysis [16]. Co-expression pertains to identification of genes which display common patterns of regulation, and may therefore be subject to similar gene regulatory mechanisms (e.g. transcription factors). Methods for co-expression analysis range from simple models of linear regression between expression values of genes [17], to construction of weighted co-expression networks consisting of gene modules [18] and deep learning-based approaches [19]. Assigning functional and biological significance to an individual gene based on co-expression requires further analysis, however, the dissection and stratification of results of co-expression analyses can be challenging [20]. This means that potentially interesting insight into functions of individual genes may be lost during transitions between methods.

Methods aiming to determine the functions of individual genes are available, and implement different approaches (see **Table 1**). Some have the objective to identify genes or genetic variation relevant to certain tissues, cell types, or cell lines (e.g. *CONTENT* [21], and *ContNeXt* [22]). While these methods are useful for the identification of significant gene-context associations, they do not predict the biological function of the given gene. Other methods use network properties (e.g. *NetDecoder* [23]) or apply coessentiality analyses (*FIREWORKS* [24]) to characterize gene-gene associations in a given context. These tools help to identify other genes significantly associated with a GOI in a context-specific manner, but again do not link these results with biological meaning. *GeneWalk* [25], *DAVID* [14, 26] and *Correlation AnalyzeR* [27] are three tools which come closest to determining the function of individual genes, in that they aim to provide context-specific biological meaning whilst being able to focus on individual genes.

**Table 1.**
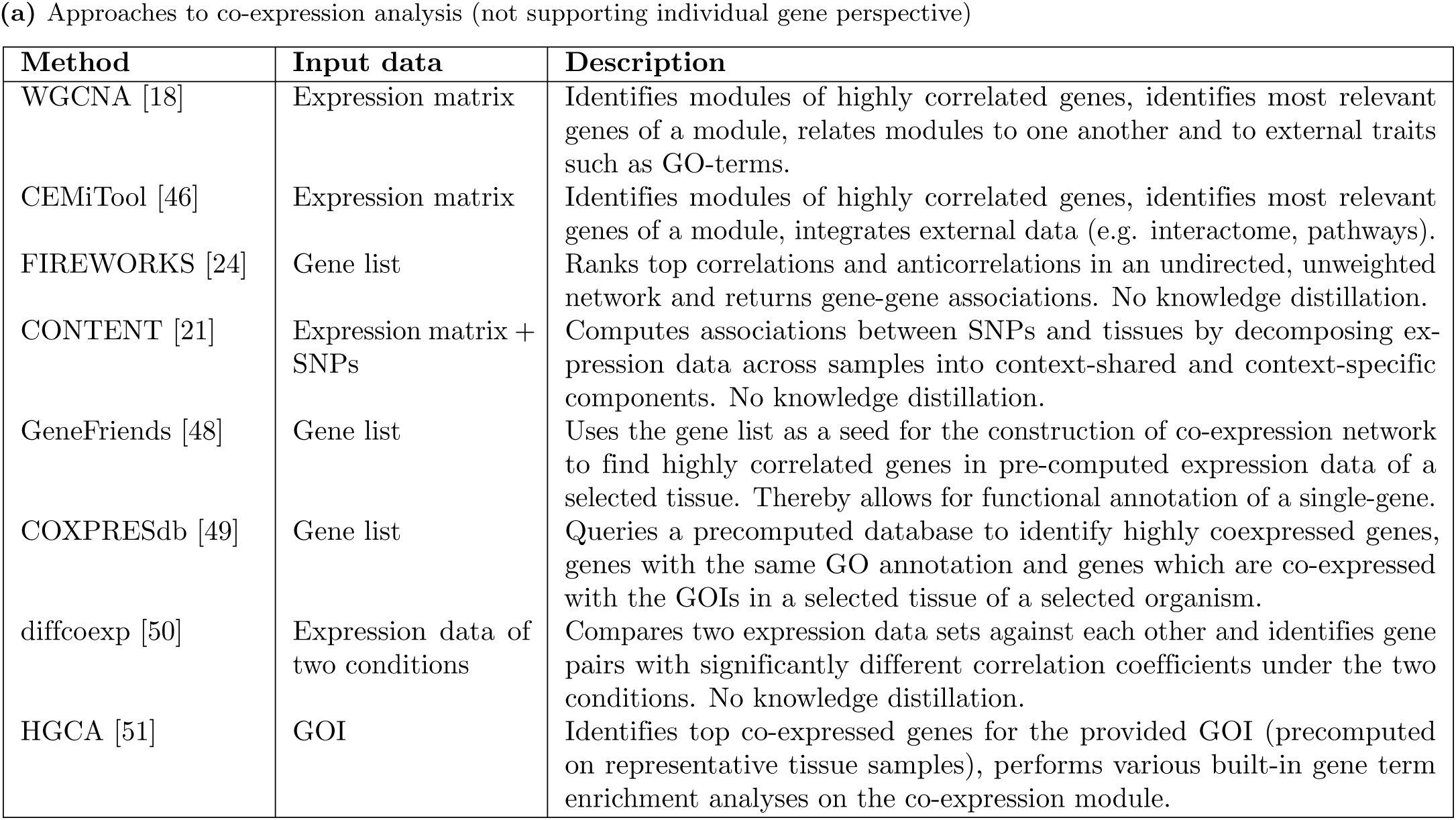
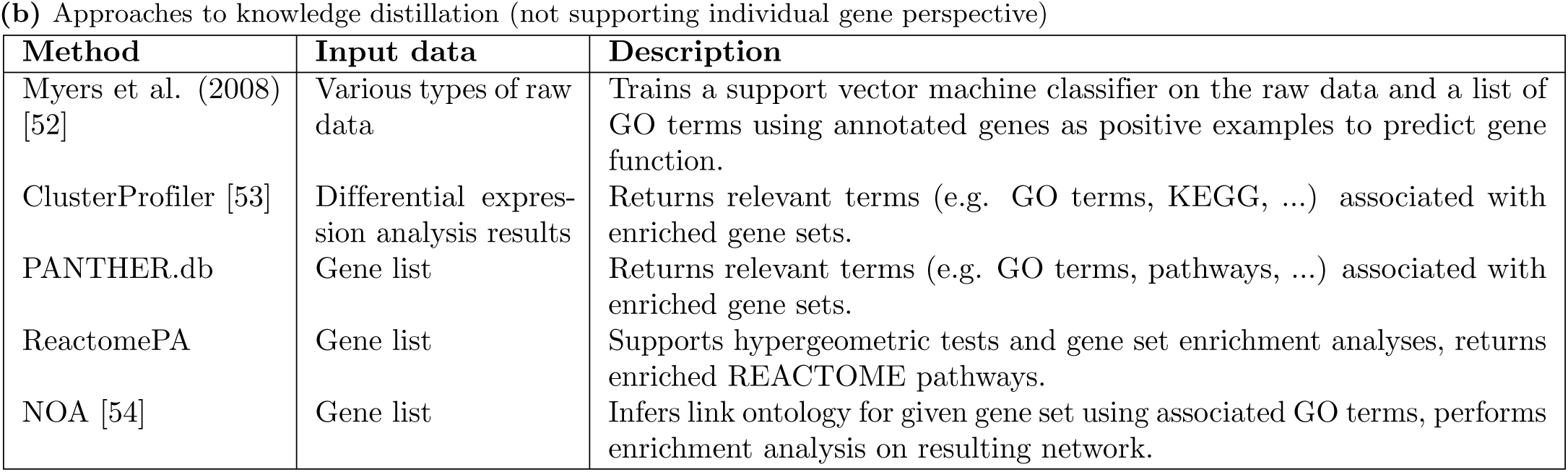
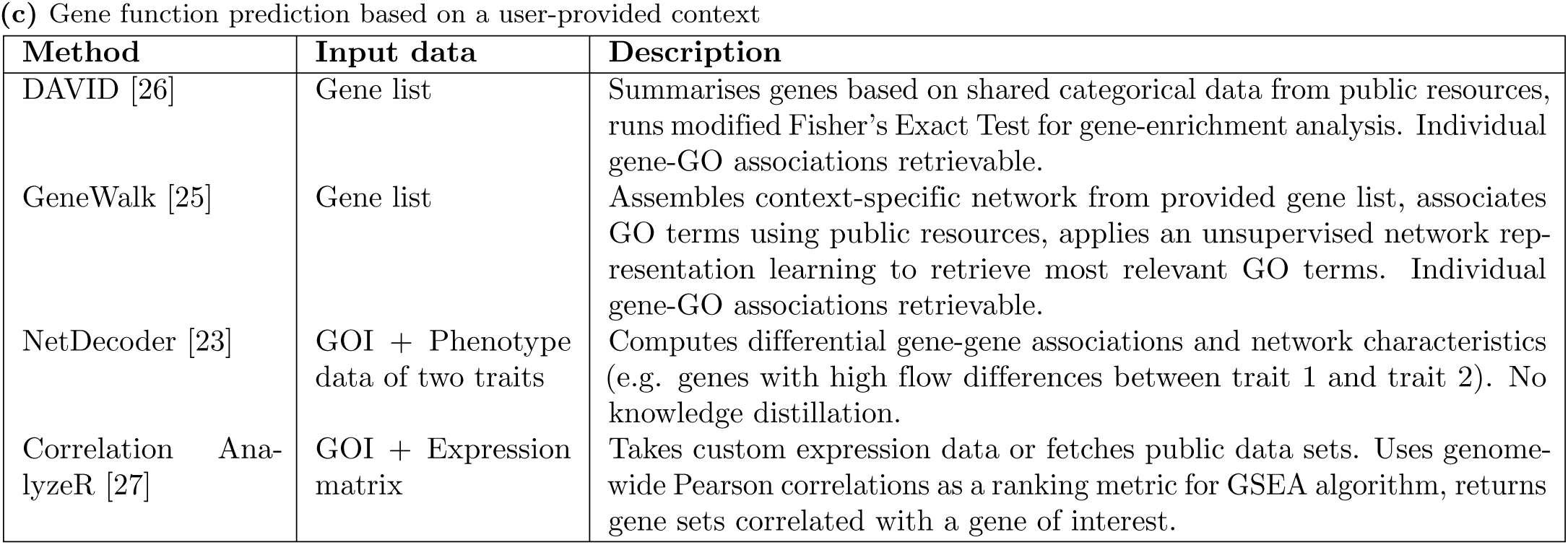
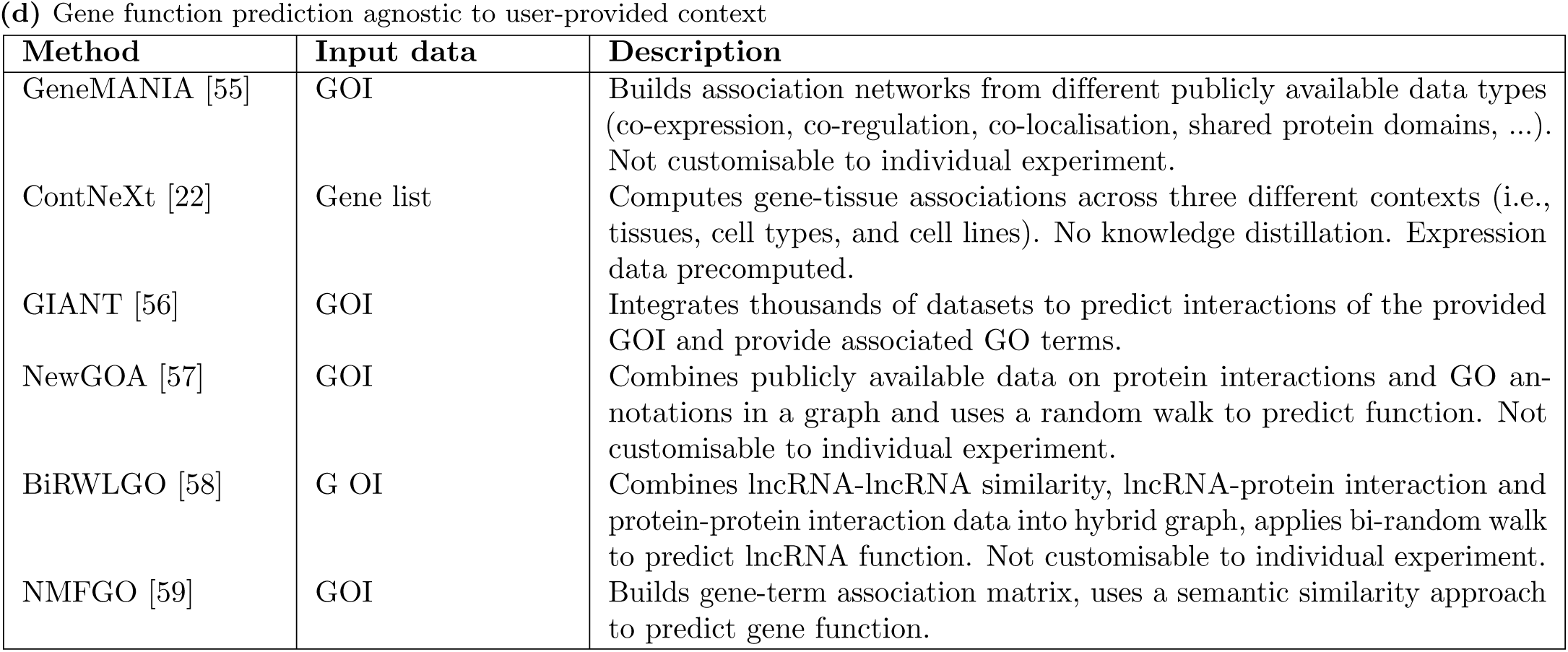
Available methods for downstream analysis of disease contexts and genes of interest.

**(a)** Approaches to co-expression analysis (not supporting individual gene perspective)
| Method | Input data | Description |
| --- | --- | --- |
| WGCNA [18] | Expression matrix | Identifies modules of highly correlated genes, identifies most relevant genes of a module, relates modules to one another and to external traits such as GO-terms. |
| CEMiTool [46] | Expression matrix | Identifies modules of highly correlated genes, identifies most relevant genes of a module, integrates external data (e.g. interactome, pathways). |
| FIREWORKS [24] | Gene list | Ranks top correlations and anticorrelations in an undirected, unweighted network and returns gene-gene associations. No knowledge distillation. |
| CONTENT [21] | Expression matrix + SNPs | Computes associations between SNPs and tissues by decomposing expression data across samples into context-shared and context-specific components. No knowledge distillation. |
| GeneFriends [48] | Gene list | Uses the gene list as a seed for the construction of co-expression network to find highly correlated genes in pre-computed expression data of a selected tissue. Thereby allows for functional annotation of a single-gene. |
| COXPRESdb [49] | Gene list | Queries a precomputed database to identify highly coexpressed genes, genes with the same GO annotation and genes which are co-expressed with the GOIs in a selected tissue of a selected organism. |
| diffcoexp [50] | Expression data of two conditions | Compares two expression data sets against each other and identifies gene pairs with significantly different correlation coefficients under the two conditions. No knowledge distillation. |
| HGCA [51] | GOI | Identifies top co-expressed genes for the provided GOI (precomputed on representative tissue samples), performs various built-in gene term enrichment analyses on the co-expression module. |

**(b)** Approaches to knowledge distillation (not supporting individual gene perspective)
| Method | Input data | Description |
| --- | --- | --- |
| Myers et al. (2008) [52] | Various types of raw data | Trains a support vector machine classifier on the raw data and a list of GO terms using annotated genes as positive examples to predict gene function. |
| ClusterProfiler [53] | Differential expression analysis results | Returns relevant terms (e.g. GO terms, KEGG, ...) associated with enriched gene sets. |
| PANTHER.db | Gene list | Returns relevant terms (e.g. GO terms, pathways, ...) associated with enriched gene sets. |
| ReactomePA | Gene list | Supports hypergeometric tests and gene set enrichment analyses, returns enriched REACTOME pathways. |
| NOA [54] | Gene list | Infers link ontology for given gene set using associated GO terms, performs enrichment analysis on resulting network. |
| DAVID [26] | Gene list | Summarises genes based on shared categorical data from public resources, runs modified Fisher's Exact Test for gene-enrichment analysis. Individual gene-GO associations retrievable. |
| GeneWalk [25] | Gene list | Assembles context-specific network from provided gene list, associates GO terms using public resources, applies an unsupervised network representation learning to retrieve most relevant GO terms. Individual gene-GO associations retrievable. |
| NetDecoder [23] | GOI + Phenotype data of two traits | Computes differential gene-gene associations and network characteristics (e.g. genes with high flow differences between trait 1 and trait 2). No knowledge distillation. |
| Correlation AnalyzeR [27] | GOI + Expression matrix | Takes custom expression data or fetches public data sets. Uses genome-wide Pearson correlations as a ranking metric for GSEA algorithm, returns gene sets correlated with a gene of interest. |

(d) Gene function prediction agnostic to user-provided context
| Method | Input data | Description |
| --- | --- | --- |
| GeneMANIA [55] | GOI | Builds association networks from different publicly available data types (co-expression, co-regulation, co-localisation, shared protein domains, ...). Not customisable to individual experiment. |
| ContNeXt [22] | Gene list | Computes gene-tissue associations across three different contexts (i.e., tissues, cell types, and cell lines). No knowledge distillation. Expression data precomputed. |
| GIANT [56] | GOI | Integrates thousands of datasets to predict interactions of the provided GOI and provide associated GO terms. |
| NewGOA [57] | GOI | Combines publicly available data on protein interactions and GO annotations in a graph and uses a random walk to predict function. Not customisable to individual experiment. |
| BiRWLGO [58] | G OI | Combines lncRNA-lncRNA similarity, lncRNA-protein interaction and protein-protein interaction data into hybrid graph, applies bi-random walk to predict lncRNA function. Not customisable to individual experiment. |
| NMFGO [59] | GOI | Builds gene-term association matrix, uses a semantic similarity approach to predict gene function. |

*GeneWalk* [25] takes a user-provided input list of genes and assembles a network composed of these genes and associated Gene Ontology (GO) terms. Network representation learning with random walks is then performed on the network. Statistical association between a given gene and GO terms is determined through comparison of node similarities between the true network and a null distribution based on node similarities in randomized networks.

Alternatively, associations between individual genes and biological functions can be performed using *DAVID* [14, 26], which takes a list of genes as input and returns GO terms, protein domain information and curated pathways which are statistically enriched in their association with a given gene, computed using Fisher’s exact test. While these approaches do provide insight into putative functions of individual genes, neither method considers the expression of the provided genes or other genes relevant to the GO terms in question. Not considering expression as a feature in these analyses could result in missing dynamic relationships between the gene-of-interest and the genes, or subsets of genes, associated with the given term. Additionally, the implementation of *GeneWalk* is limited to the use of GO terms, and cannot be implemented with other curated gene sets which may provide more relevant functional annotations in a specific context, such as disease.

One method which considers co-expression and outputs putative gene function is *Correlation AnalyzeR* [27]. Here, weighted Pearson correlations between normalized gene expression counts are calculated between a gene-of-interest and other genes present in the expression data. A ranked gene list is then assembled from the resulting correlation values, which is used as input to gene set enrichment analysis, resulting in statistically enriched terms which are theoretically co-expressed with the gene-of-interest. However, the authors state that for a robust analysis, datasets of more than 30 samples and at least 4 different studies should be used, limiting the contexts in which this method can be used.

We sought to explore how co-expression and functional enrichment analyses can be combined into a single workflow which provides insight into the function of a specific GOI in a given context provided by the input data. Such a method would permit a comprehensive assessment of expression patterns and putative functions of a GOI across multiple experimental conditions using experimental data generated by the user. To this end, we propose *GeneCOCOA*, an *R* package which identifies and ranks functional gene sets which are co-expressed with a user-supplied GOI. *GeneCOCOA* may be run using either user-supplied or publicly available gene expression data, and can utilize several curated databases of gene annotations in order to compute functional enrichments in co-expression.

## Materials and methods

### Databases

For the functionality of GeneCOCOA described herein, curated gene sets from the Hallmark database [28], as well as genes annotated to the Biological Process domain of Gene Ontology [29] (GO:BP) were used.

### Input data

The use cases described in this manuscript utilized publicly available transcriptome profiling data available from *Gene Expression Omnibus* [2,3] under the accession numbers GSE36980 [30], GSE28253 [31], GSE5406 [32], GSE9006 [33], GSE48060 [34], GSE17048 [35], and GSE114922 [36].

The RNA-sequencing data arising from acute myeloid leukemia patients [37] is available publicly from the *European Genome-Phenome Archive* [38] under the accession EGAD00001008484, and initial access prior to the publication of the data was provided by Prof. Dr. Thomas Oellerich and Dr. Sebastian Wolf (Goethe University Frankfurt, University Hospital Frankfurt).

### Preprocessing

Raw reads were aligned against the *hg38* genome using *Bowtie2* (v2.3.5.1) [39], with default parameters, and quantified using *Salmon* (v1.5.2) [40], with default parameters. Curated quantified and normalized expression data sets were fetched with *gemma.R* [41].

### Detection of gene sets which are co-expressed with a gene-of-interest

#### Determining number of gene subsets

The number of gene subsets sampled from each gene set *i* is implemented as a user-controlled parameter. In test runs, we determined *i* = 1000 to provide an acceptable compromise between efficiency and statistical power (see **Supplementary Figure S1**). Therefore, we set *i* = 1000 for all analyses in this manuscript.

#### Generation of gene subsets

Initially, a number of subsets (default 1000) are derived from a given gene set (e.g. pathway, GO term), as described by the following:

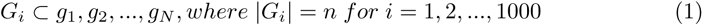

where *G_i_* is the *i*-th subset of *n* genes *g*_1_*, g*_2_*, …, g_N_* which make up the total gene set *G*.

#### Linear regression models

The dataset-specific expression values of each gene in a subset of genes serve as predictor variables in a linear regression model with the expression of a GOI being the outcome variable, as described by:

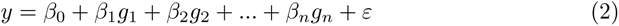

where *g*_1_*, g*_2_*, …, g_n_* represent the dataset-specific expression values of the genes of the subset, *β*_0_*, β*_1_*, β*_2_*, …, β_n_* are the coefficients for each predictor variable, *y* represents the predicted expression of the GOI, and *ε* represents the error of the linear regression model.

#### Root-mean-square error calculation

For each gene subset, the linear regression model produces predicted values *ŷ_i_*based on the predictors *g_i_*. The root-mean-square error (RMSE) for the *i*-th subset is then calculated as:

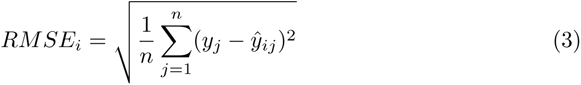

where *y_j_*is the true expression of the GOI and *y*^*_ij_* is the predicted expression from the linear regression model for the *j*-th observation using the subset *G_i_*.

The same procedure is performed for a size-matched set of randomly sampled genes, resulting in two sets of RMSE values. One derived from linear regression models predicting the expression of the GOI from subsets of genes from a given gene set, and one derived from linear regression models predicting the expression of the GOI from randomly sampled subsets of genes expressed in the given dataset.

#### Computation of gene set-specific enrichment P values

RMSE values *RMSE_i_, …, RMSE*_1000_ derived from subsets *G_i_, …, G*_1000_ of a given gene set *G* are compared against RMSE values *_r_RMSE_i_, …, _r_RMSE*_1000_ derived from randomly subset genes using a Student’s t-test. The resulting *P* value is subsequently adjusted using the Benjamini-Hochberg method [42]. This results in an adjusted *p* value for each gene set in a given curated database, describing the strength of association between the genes comprising each gene set and the user-provided GOI.

#### Inter-gene set comparison and visualization

In order to further stratify gene sets of potential interest, the direction of co-expression between the GOI and curated gene sets is also calculated during the course of the workflow, and is included as a parameter for visualization of results. This value is calculated as follows:

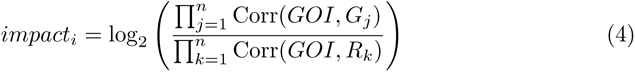

where Corr(*GOI, G_j_*) represents the correlation between the GOI and a gene of a given gene set *G* of size *n*, and Corr(*GOI, R_k_*) represents the correlation between the GOI and a gene from a size-matched set of randomly sampled genes *R*. *impact_i_ >* 0 thus indicates that the co-expression between the GOI and the gene set *G* is smaller than a random baseline mean co-expression.

#### Comparison to similar methods

GeneCOCOA was compared to other methods which aim to annotate the functions of individual genes by testing the ability of each tool to accurately link genes associated with a given disease to GO terms implicated in the same disease. The methods considered for comparison were DAVID [14], GeneWalk [25] and Correlation AnalyzeR [27].

#### Definition of disease-relevant genes

In order to define a relevant gene set for each condition to be studied, the DisGeNET [43] platform was queried via web interface (https://www.disgenet.org/search) with the full name of each condition (“Amyotrophic Lateral Sclerosis”, “Alzheimer’s Disease”, “Dilated Cardiomyopathy”, “Insulin-dependent Diabetes Mellitus”, “Myocardial Infarction” and “Multiple Sclerosis”). As of 14-06-2023, the top-ranked hits were the entries with the UMLS/concept IDs C0002736, C0002395, C0007193, C0011854, C0027051 and C0026769. From each summary of gene-disease associations (GDA), genes with a *Score_gda_* ≥ 0.5 were considered as substantially associated with the disease and included in the input set of disease-relevant genes.

#### Definition of disease-relevant terms/gene sets

To obtain disease-relevant gene sets, the MalaCards database [44] was queried via web interface (https://www.malacards.org/) with the full name of the condition (“Alzheimer’s Disease”, “Amyotrophic Lateral Sclerosis”, “Dilated Cardiomyopathy”, “Insulin-dependent Diabetes Mellitus”, “Myocardial Infarction” and “Multiple Sclerosis”) on 14-06-2023. The top hit was selected based on the MalaCards InFormaTion Score and the Solr relevance score provided by MalaCards. For each disease card (MalaCards IDs ALZ065, AMY091, DLT002, TYP008, MYC007 and MLT020, respectively), the complete list of Gene Ontology Biological Process terms was downloaded and treated as the ground truth collection *T* for the respective disease.

#### Construction of input gene lists for GeneWalk and DAVID

To assemble a context-specific gene network, GeneWalk requires a list of relevant genes obtained from a specific experimental assay as an input. To this end, GEO2R [3] was used to obtain a list of differentially expressed (DE) genes for each of the publicly available transcriptomic gene sets. Any gene with an adjusted *p <* 0.05 between control and disease condition, as calculated by *DESeq2* [45], was considered differentially expressed. To ensure that all disease-relevant genes obtained via DisGeNET would be included as well, the union of disease-relevant genes and DE genes was obtained. Thus, a context-set *C* was created for each condition.

### Systematic comparison of GeneCOCOA, Correlation AnalyzeR, GeneWalk and DAVID

Each method was used to determine the association of disease-relevant genes (as per defined via DisGeNET, see subsection *Definition of disease-relevant genes*) with disease-relevant gene sets (as defined via MalaCards, see previous subsection). Since GeneWalk results are computed on Gene Ontology annotations [7], we restricted the comparison to gene sets from the GO:BP collection.

The ability of each tool to report any disease-relevant GO:BP term for a list of disease-relevant genes-of-interest across different diseases was tested. We distinguished two cases: (1) A disease-relevant gene is analyzed in a condition matching its disease. In this case, we expect the method to report a significant association between the gene and any of the disease-relevant GO:BP terms reported in MalaCards (”true positive”). (2) A disease-relevant gene-of-interest is analyzed using data arising from a separate disease where said gene is not annotated as being important in DisGeNet, therefore a significant association between the gene and the terms present in the MalaCard for the disease is not expected (”false positive”).

*GeneCOCOA*, *GeneWalk*, *DAVID* and *Correlation AnalyzeR* [27] were run for every combination of disease-relevant genes – Alzheimer’s Disease (AD): 24, Amyotrophic Lateral Sclerosis (ALS): 16, Dilated Cardiomyopathy (DC): 12, Diabetes Mellitus (DM): 4, Myocardial Infarction (MI): 21, Multiple Sclerosis (MS): 7 – and diseases. In each case, a disease-specific expression data set was provided as input, a single disease-relevant gene was provided as the gene-of-interest, and the GO:BP ontology provided as the collection of gene sets to rank. For each disease, *GeneWalk* and *DAVID* were run with the appropriate context-set *C* (see previous subsection) as the input list (including the additional genes-of-interest which are not functionally linked to the disease in question, see case (2) above). Gene-of-interest-associated GO:BP terms were parsed from the results of each method using a threshold of *p_adjusted_ <* 0.05.

### GeneCOCOA R package

*GeneCOCOA* is formulated as an R package and is hosted on GitHub at the URL https://github.com/si-ze/geneCOCOA.

## Results

### *GeneCOCOA* identifies functional gene sets co-expressed with a gene-of-interest

The COmparative CO-expression Analysis focused on a Gene-of-interest (*GeneCOCOA*) presented here incorporates multiple approaches which aim to functionally annotate genes following gene expression profiling (**Fig. 1A**). Several approaches exist for the analysis of experiment-specific co-expression patterns (e.g. *WGCNA* [18], *CemiTool* [46]), the harnessing of curated knowledge (e.g. Molecular Signature Database (*MSigDB* ) [28]), as well as for the integration of prior knowledge with experiment-specific co-expression patterns (e.g. *GSEA* [47], *Enrichr* [12]). Some methods also aim to apply prior knowledge to predict the functions of individual genes, most notably *DAVID* [26], *GeneWalk* [25] and *Correlation AnalyzeR* [27]. However, few methods exist which utilize co-expression and curated gene sets to predict gene function (summarized in **Table 1**). To our knowledge, only *Correlation AnalyzeR* [27] provides this option in *single-gene mode*. Yet, its results are based on a single correlation analysis. *GeneCOCOA* has been developed as an integrative method which aims to apply curated knowledge to experiment-specific expression data in a gene-centric manner based on a robust bootstrapping approach.

**Figure 1.**
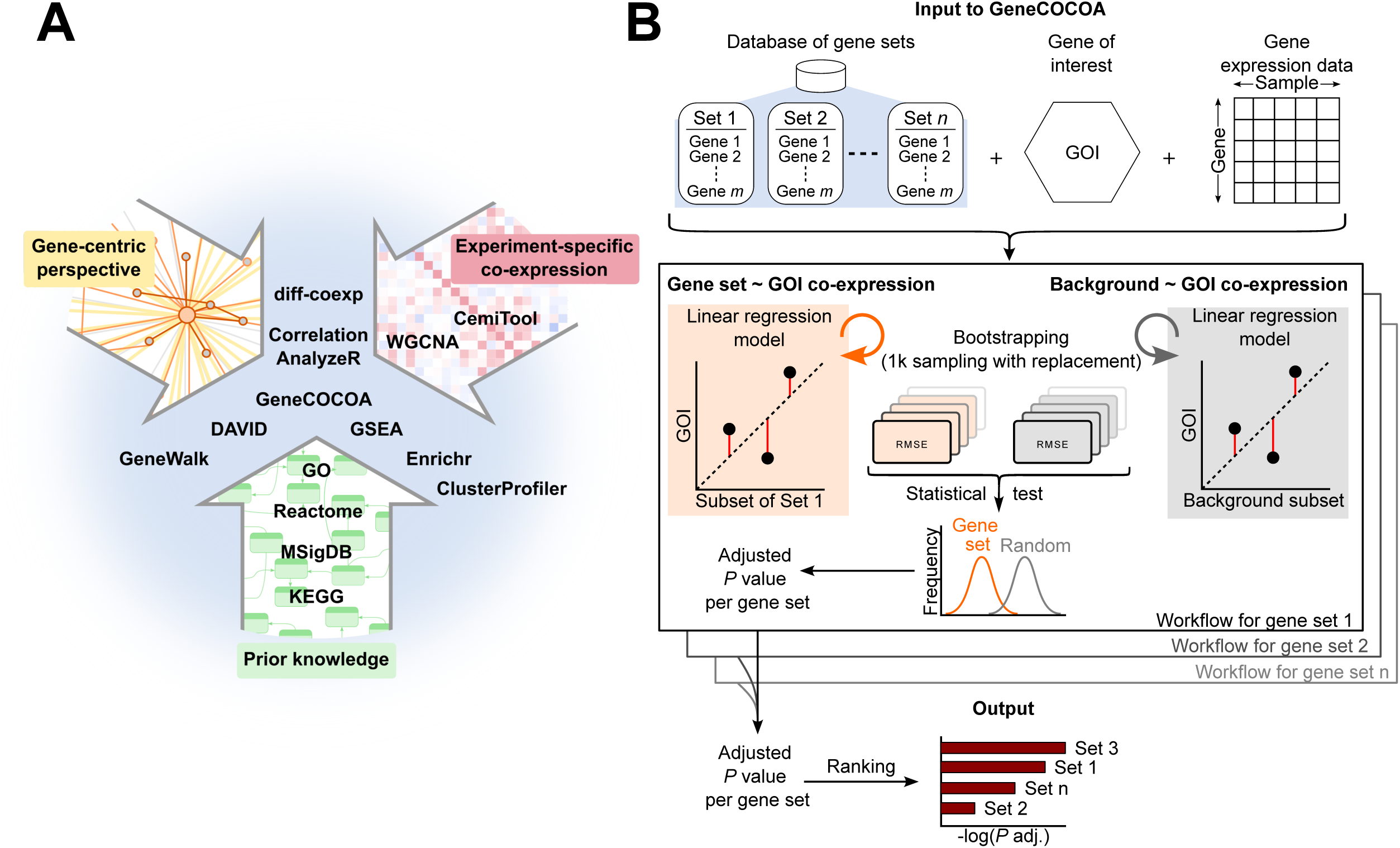
GeneCOCOA workflow for identification of functional gene sets co-expressed with a gene-of-interest. **(A)** Strategies and related methods for statistically associating genes to putative functions, summarized into gene-centric (GeneWalk, DAVID), prior knowledge (GO, Reactome, MSigDB) and co-expression (WGCNA, CemiTool) approaches. GeneCOCOA incorporates elements of each of these approaches into a single workflow. **(B)** Schematic representation of the GeneCOCOA workflow, which takes as input user-provided functional gene sets, a gene-of-interest (GOI) and gene expression data to report statistically ranked gene sets associated with the provided GOI. This is achieved by comparing root-mean-square error (RMSE) values from bootstrapped linear regression models predicting the expression of the GOI using either genes arising from a single gene set, or randomly sampled genes from the expression data. Gene set errors and random errors are statistically compared, and the resulting *p* values are adjusted, resulting in an output list of functional gene sets ranked statistically by the strength of their association with the provided gene-of-interest.

The input to *GeneCOCOA* is a list of curated gene sets (e.g. from Gene Ontology, MSigDB, pathways), a gene-of-interest (GOI) that the user wishes to interrogate, and a gene expression matrix of *sample ∗ gene* (**Fig. 1B, top**). From each gene set, *n* genes are sampled and used as predictor variables in a linear regression modelling the expression of the GOI as the outcome variable (**Fig. 1B, middle**). A background model is created analogously by sampling *n* random genes from the complete expression data set. For bootstrapping, this procedure is repeated *i* times, *i* being a parameter that can be specified by the user. Testing different values of *i*, we found *i* = 1000 to provide the best tradeoff between efficiency and power (see **Supp. Fig. S1**). The *i* gene set model errors and *i* random model errors are compared in a t-test. Gene sets with *p_adjusted_ <* 0.05 are considered to model the expression of the GOI better than random, and the *p_adjusted_* values are used to stratify and rank gene sets (**Fig. 1B, bottom**). The results output by *GeneCOCOA* aim to provide insight into potential functions of the gene-of-interest in the specific context provided by the gene expression data.

### Detection of context-specific changes in gene function using *GeneCOCOA*

To test the ability of *GeneCOCOA* to detect changes in gene function resulting from disease, it was applied to identify functions of the gene FMS-like tyrosine kinase 3 (*FLT3* ) in acute myeloid leukemia (AML). AML is a malignancy of the hematopoietic system affecting the differentiation and maturation of myeloid blood cells. Characterized by a complex genetic landscape, AML can be divided into various subtypes, which differ in both phenotype and prognosis. One common (25% of patients [60]) mutation linked to AML is the internal tandem duplication (ITD) of *FLT3*. Normally, expression and activation levels of *FLT3* are important for maintaining a balance of proliferation and differentiation in hematopoietic cells [61]. *FLT3* -ITD results in a constitutive activation of the kinase, promoting a hyperproliferative state and cell survival [62]. *FLT3* -ITD is associated with a higher disease burden, higher relapse rate and inferior overall survival [14].

A whole-transcriptome RNA-sequencing dataset of 136 AML patients [37] was subset for patients with *FLT3* -ITD mutations (31 patients). Taking *FLT3* as the GOI, *GeneCOCOA* was used to assess the significance of the association between *FLT3* and gene sets defined by GO Biological Processes (GO:BP). For comparison, a control set of 48 healthy *CD34+* bone marrow samples was constructed from data under the GEO accession GSE114922 [36]. Again, *GeneCOCOA* was used to detect and rank associations between *FLT3* and GO:BP terms (**Fig. 2A**).

**Figure 2.**
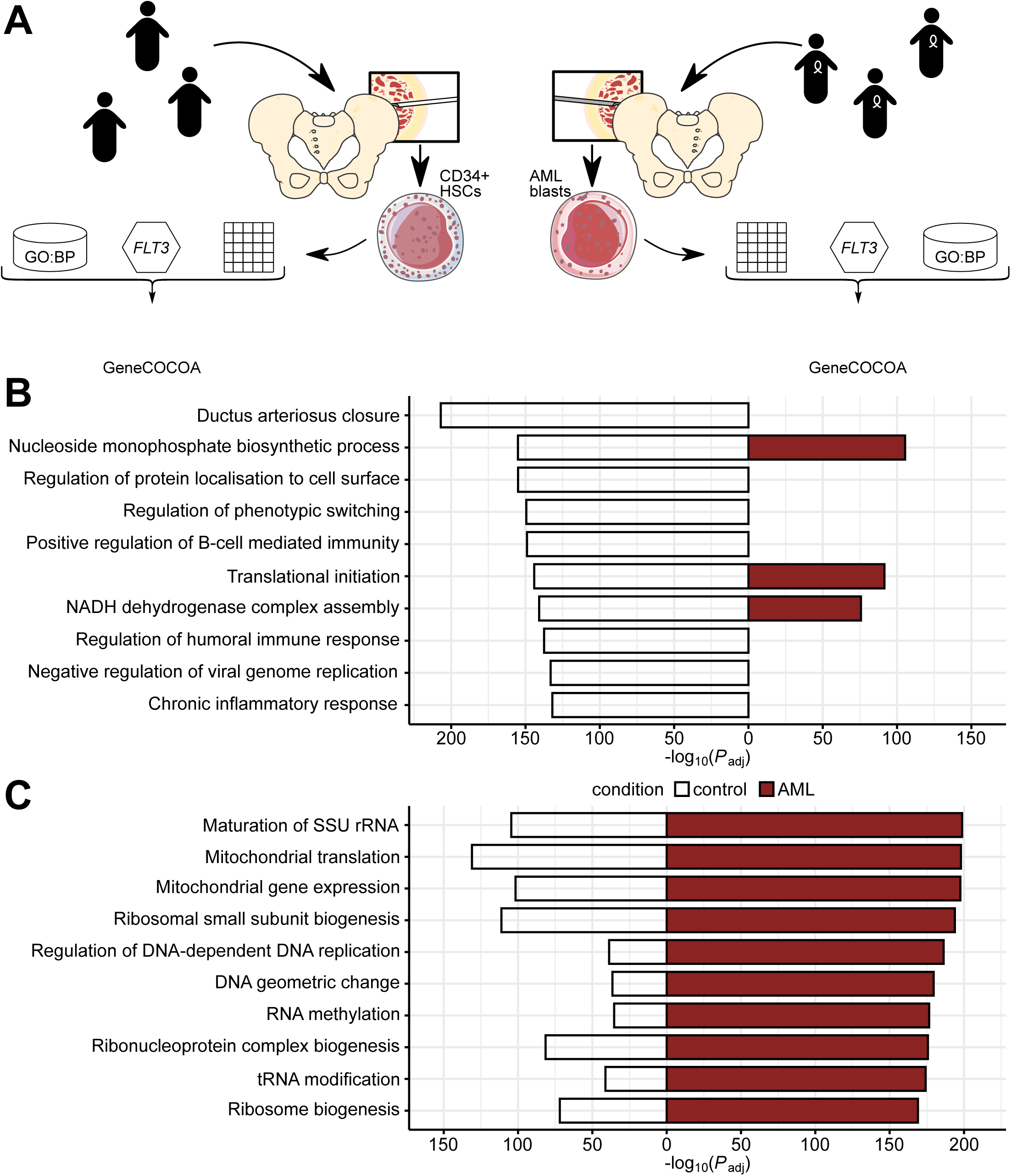
Example use case of GeneCOCOA to predict context-specific FLT3 function using expression data from hematopoietic stem cells and acute myeloid leukemia blasts. **(A)** In an exemplary use case, GeneCOCOA was applied to study the co-expression patterns of *FLT3* with Gene Ontology Biological Process (GO:BP) terms in bulk RNA-sequencing of CD34+ hematopoietic stem cells (HSCs) from 48 healthy subjects, and blasts from 31 patients with acute myeloid leukemia (AML) positive for *FLT3*-ITD mutations. **(B)** The 10 highest-ranked GO:BP terms with *FLT3* in HSCs from healthy donors, as computed by GeneCOCOA. The corresponding significance values in AML blasts are provided for comparison. Ranks are annotated next to the bars; non-significant terms are not annotated. **(C)** The 10 highest-ranked GO:BP terms with *FLT3* in patients with AML and *FLT3*-ITD mutations, as computed by GeneCOCOA. The corresponding significance values in healthy HSCs are provided for comparison. Ranks are annotated next to the bars; non-significant terms are not annotated.

Physiologically, *FLT3* is involved in immune function and regulation of hematopoietic cell proliferation and differentiation [61]. Accordingly, among the GO:BP terms associated with *FLT3* by *GeneCOCOA* in healthy *CD34+* cells are terms associated with immune response (e.g. ”Regulation of humoral immune response”, ”Chronic inflammatory response”) and terms indicating both proliferative processes (e.g. ”Nucleoside monophosphate biosynthetic process”) and differentiation (e.g. ”Positive regulation of B-cell mediated immunity”, ”Regulation of phenotypic switching”) (**Fig. 2B**). This complex profile is lost in the *GeneCOCOA* results for *FLT3* co-expression patterns in AML blasts (**Fig. 2C**). The top 10 GO:BP terms reflect mitochondrial processes (e.g. ”Mitochondrial gene expression”) and cell growth/division (e.g. ”Regulation of DNA-dependent DNA replication”, ”Ribosome biogenesis”), reflecting the switch to a predominantly proliferative profile. The results thus replicate dysregulation of *FLT3* expression and function previously described in literature, indicating that *GeneCOCOA* may be able to detect context-dependent changes in gene function, given appropriate data.

### *GeneCOCOA* detects disease-driven alterations in gene co-expressio patterns

In further proof-of-principle testing, *GeneCOCOA* was applied to gene expression datasets arising from diseases with well-studied causative genes. Here, the direction of co-expression between the gene-of-interest and each curated gene set was also analyzed in each case (denoted ’*Impact* ’, see **Eq. 4** in *Materials and methods*). Disease data sets were compared against healthy control data sets in terms of their co-expression patterns between a known causative GOI and fifty MSigDB Hallmark gene sets [28].

One disease in which causative genes have been suggested in literature is amyotrophic lateral sclerosis (ALS). The first gene to be identified as causative for this neurodegenerative disease was superoxide dismutase 1 (*SOD1* ) [63]. *SOD1* codes for Cu/Zn superoxide dismutase type-1, an enzyme crucial for cellular antioxidant defense mechanisms. Mutations of *SOD1* in ALS are known to destabilize the protein, leading to misfolding. This triggers various pathophysiological events such as protein accumulation, mitochondrial and/or proteasome dysfunction and accumulation of reactive oxygen species (ROS). This switch between contexts is reflected in the *GeneCOCOA* results when *SOD1* is taken as the gene-of-interest, along with gene expression data from disease and healthy conditions. When *GeneCOCOA* was run with gene expression data from lymphocytes of 11 healthy donors, *SOD1* was closely linked with immune function (e.g. ”Allograft rejection”, ”TNF-*α* signalling via NF-*κ*B”, ”Inflammatory response”), as well as gene sets related to oxidative stress (e.g. ”Peroxisome”, ”ROS Pathway”) (**Fig. 3A, left**). In accordance with literature [64, 65], many of these associations are lost in the lymphocyte transcriptomes of 11 patients with ALS. Instead, a gain in association between *SOD1* and genes associated with oxidative phosphorylation could be observed, reflecting potential mitochondrial defects (**Fig. 3A, right**). Also indicative of the pathophysiology of *SOD1* -driven ALS was the association between *SOD1* expression and the Hallmark gene set ”Unfolded protein response”. The detection of this term – specifically in the disease samples – demonstrates that *GeneCOCOA* has the potential to identify context-specific co-expression patterns with disease relevance.

**Figure 3.**
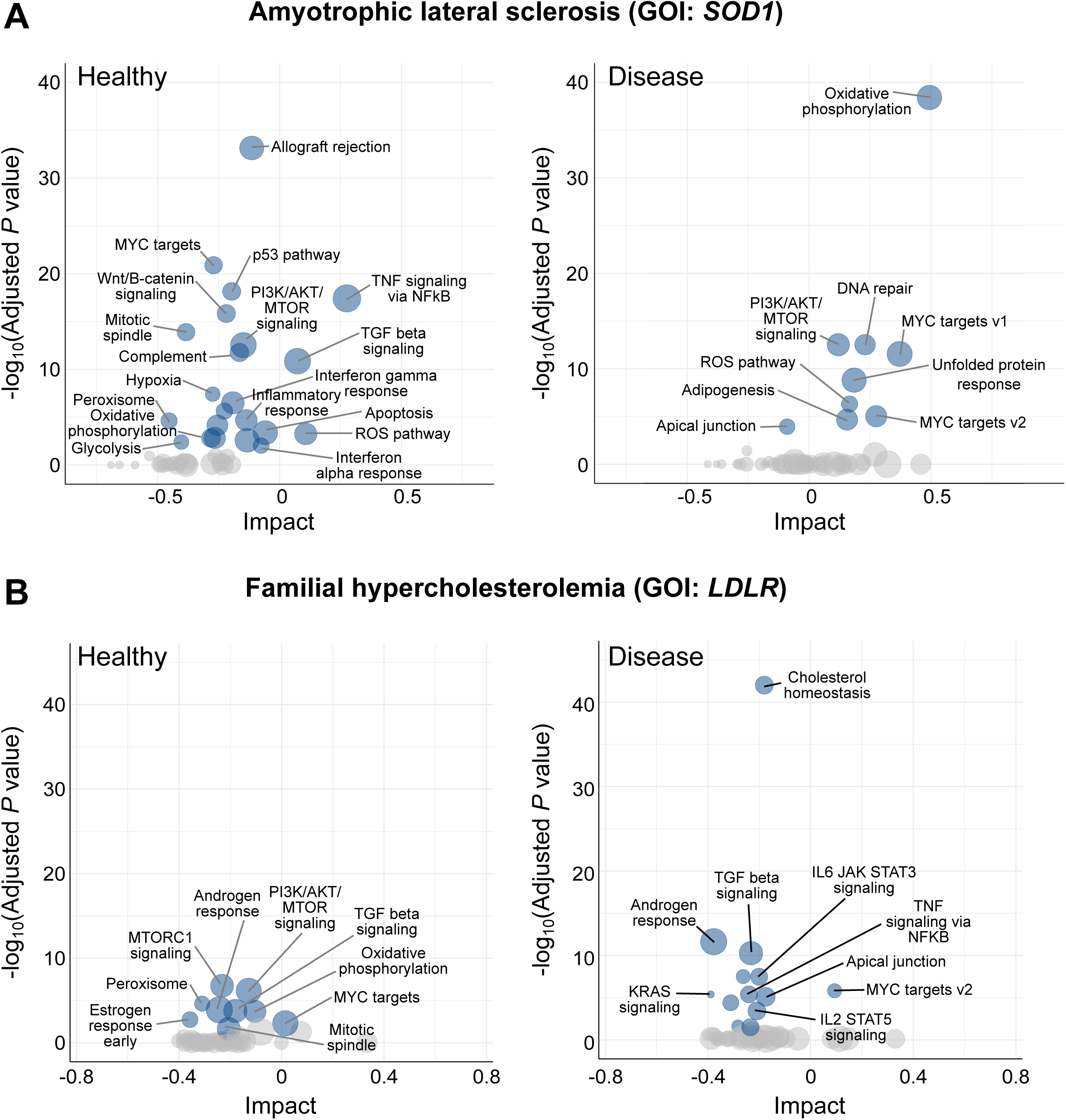
Gene-COCOA detects cellular responses to diseases with monogenic signatures. **(A)** GeneCOCOA results reporting the strength of association in co-expression between *SOD1* and MSigDB Hallmark gene sets, in lymphocytes isolated from healthy donors (left) and lymphocytes isolated from patients with amyotrophic lateral sclerosis (right). **(B)** GeneCOCOA results reporting the strength of association in co-expression between *LDLR* and MSigDB Hallmark gene sets in monocytes from healthy donors (control, left) and monocytes isolated from patients with familial hypercholesterolemia (disease, right). The size of the points in each plot reflects the relative mean expression level of each gene set.

In another use case, *GeneCOCOA* was run using gene expression data originating from isolated lymphocytes of 10 patients with familial hypercholesterolemia (FH), comparing them to 13 healthy control samples. FH is an autosomal dominant disorder of lipoprotein metabolism characterized by high levels of cholesterol. The most common causes are mutations in the gene coding for low-density lipoprotein receptor (*LDLR*). Physiologically, the LDL transmembrane receptor mediates the internalization and lysosomal degradation of LDL. Mutations disrupting the function of LDLR lead to elevated plasma levels of LDL, promoting accelerated atherosclerosis and coronary heart disease [66, 67]. In correspondence with these mechanisms described in literature, the *GeneCOCOA* results indicated that the functional association between *LDLR* and genes annotated to be important for ”Cholesterol homeostasis” became stronger in FH samples compared to control samples (**Fig. 3B**). Again, these results suggest that *GeneCOCOA* is able to detect changes in gene co-expression which are pertinent to disease-specific conditions. While these results were promising, the question remained of how the approach implemented in *GeneCOCOA* compared to methods with the similar aim of functionally annotating individual genes.

### *GeneCOCOA* provides a comprehensive gene-focused co-expression and functional analysis missing from similar methods

To our knowledge, only few approaches to the problem of inferring the function of a specific gene-of-interest (GOI) been published (**Table 1**), most notably *DAVID* [14], *GeneWalk* [25] and *Correlation AnalyzeR* [27].

*DAVID* is a web-accessible set of functional annotation tools which allows for the rapid mining of a wide range of public resources. Provided with a list of gene identifiers, *DAVID* summarizes them, based on shared categorical data in gene ontology, protein domain, and biochemical pathway membership, returning a modified Fisher Exact *p*-value for gene-enrichment analysis.

*GeneWalk* allows for the GO enrichment analysis of an experiment-specific gene set (e.g. differentially expressed genes). Using publicly available resources, *GeneWalk* first assembles a context-specific gene network which represents both interactions between the provided genes and links to GO terms, then applies an unsupervised network representation learning algorithm (*DeepWalk* [68]) to retrieve the GO terms of highest statistical relevance.

*Correlation AnalyzeR* [27] has been developed for the exploration of co-expression correlations in a given data set, and in *single-gene mode* also supports the prediction of individual gene functions and gene-gene relationships. In an adaption of the Gene Set Enrichment Analysis [47] (GSEA) algorithm, it employs genome-wide Pearson correlations as a ranking metric to determine the gene sets correlated with a GOI.

*GeneCOCOA* and *Correlation AnalyzeR* [27] exploit the user-provided expression data to gain insight into gene correlations in a context-specific manner. GeneWalk and, less explicitly, *DAVID*, require a list of input genes to assemble the context. Using *gemma.R* [41] and *GEO2R* [3] for the selection of potential input data sets, we therefore focused on sufficiently large (*n >* 10) transcriptomic data sets in which we could reliably identify a set of DE genes. Six curated data sets met our criteria. Disease-relevant GO:BP terms were then retrieved from *MalaCards* [44], and disease-relevant genes from *DisGeNET* [10, 43].

In a systematic comparison, *DAVID*, *GeneWalk*, *Correlation AnalyzeR* [27] and *GeneCOCOA* were used to search for statistically significant associations between matching disease-relevant genes and disease-relevant GO:BP terms (**Fig. 4A**). Each method was run for every combination of disease (AD: Alzheimer’s Disease, ALS: Amyotrophic Lateral Sclerosis, DC: Dilated Cardiomyopathy, DM: Insulin-dependent Diabetes Mellitus, MI: Myocardial Infarction and MS: Multiple Sclerosis) and disease-relevant genes (total genes AD: 24, ALS: 16, DC: 12, DM: 4, MI: 21, MS: 7). For each method, a statistically significant (*p_adjusted_ <* 0.05) association between a given gene and a condition-relevant term was recorded. If the gene belonged to the matching disease-relevant gene set, this was considered a true positive, whereas if the gene was a member of one of the other disease sets, it was considered a false positive. Although these terms are not strictly accurate given the nature of these types of analysis, they are used here in an attempt to compare these methods in an objective and unbiased manner, and this matter is further covered in the *Discussion*.

**Figure 4.**
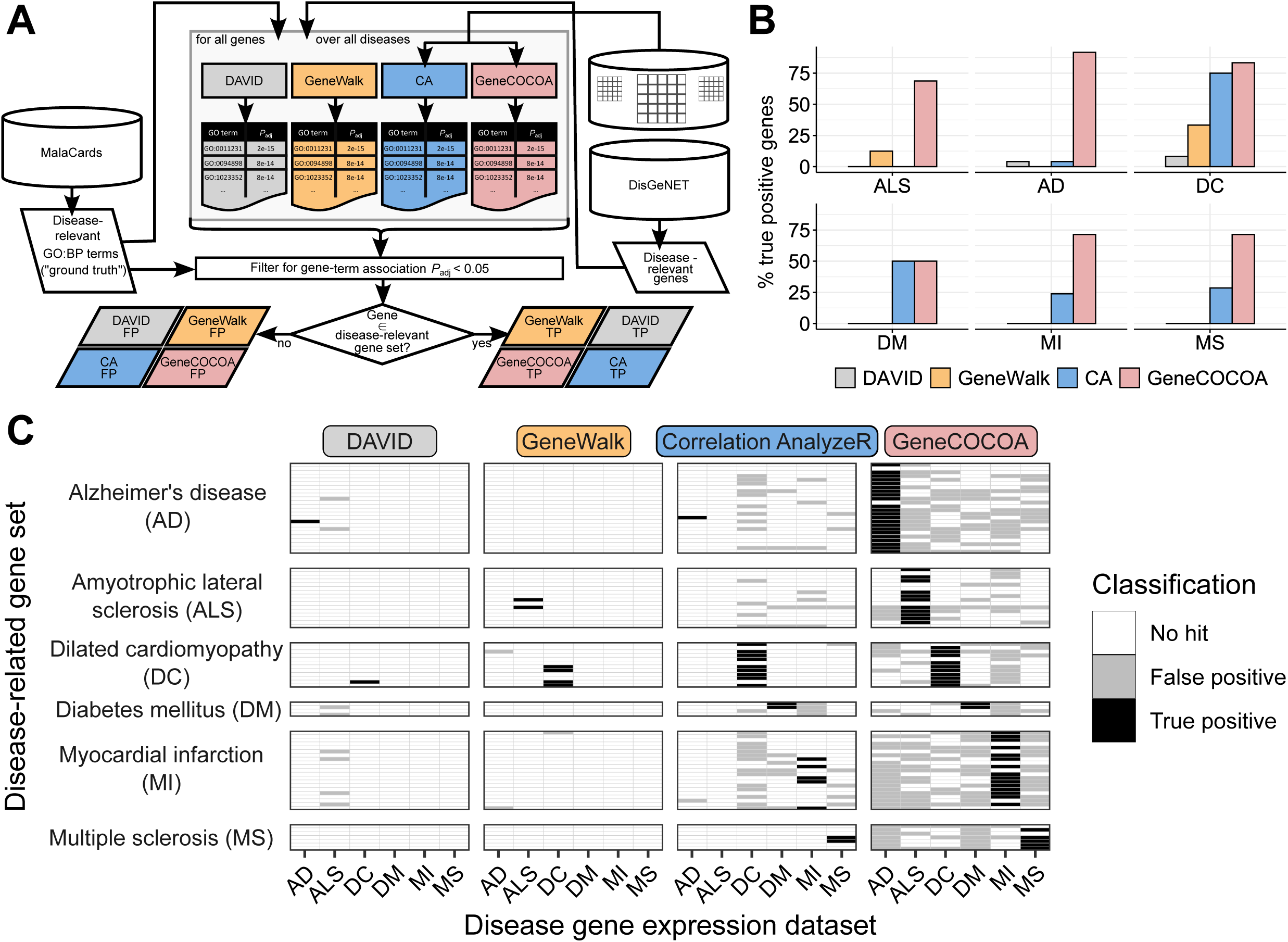
Systematic comparison of GeneCOCOA, DAVID, Correlation AnalyzeR and GeneWalk for their performance in statistically linking disease-relevant genes and GO:BP terms. **(A)** GeneCOCOA, DAVID, Correlation AnalyzeR (CA) and GeneWalk were each run to identify significantly associated disease-relevant genes from DisGeNet and disease-associated Gene Ontology Biological Process terms (GO:BP) as listed on MalaCards. Genes significantly associated to the matching disease terms were considered true positives (TP), and genes statistically linked to terms from other diseases as false positives (FP). **(B)** Proportion of true positive associations between disease-relevant genes and matching disease GO:BP terms by GeneCOCOA, GeneWalk, Correlation AnalyzeR (CA) and DAVID (AD: Alzheimer’s disease, ALS: Amyotrophic lateral sclerosis, DC: Dilated cardiomyopathy, DM: Diabetes mellitus, MI: Myocardial infarction, MS: Multiple sclerosis). **(C)** Summary of true positive and false positive gene-term associations per set of disease-relevant genes across all diseases, as computed by GeneCOCOA, GeneWalk, Correlation AnalyzeR and DAVID.

Across all conditions, *GeneCOCOA* had a substantially higher true positive rate than either *DAVID* or *GeneWalk*, and in all but one case also a higher true positive rate than *Correlation Analyzer* (**Fig. 4B**). In order to confirm that *GeneCOCOA* was not just returning spurious significant associations for every provided gene, the proportions of false positives across all conditions for all methods was further analyzed. Overall, *GeneCOCOA* reported more false positives than the other methods (**Fig. 4C**, **Supp. Fig. S2 & S3**). However, when considering the results in a gene-set-focused perspective, *GeneCOCOA* recalls more true positives per gene set than false positives (corresponds to the summary of row counts in **Fig. 4C**; see also **Supp. Fig. S2, S3 & S4**. This is truly independent of the disease expression set provided. From a condition-wise perspective (corresponding to columns in **Fig. 4C**), *GeneCOCOA* consistently reports a higher proportion of true positives than false positives across all conditions (**Supplementary Figure S3**). For *GeneWalk* and *DAVID*, the proportions of true and false positives were negligible, resulting in both methods having high true negative rates, but accompanying high false negative rates as well. *Correlation Analyzer* managed to recover more true positives than the prior two methods, yet in the majority of cases the false positive rate was at least as high as the true positive rate (see **Supp. Fig. S3** ). Thus, GeneCOCOA recovers the most relevant disease terms whilst maintaining an acceptable level of specificity, independent of disease type.

Taken together, the results presented here demonstrate that *GeneCOCOA* is capable of identifying statistically significant functional co-expression patterns linked to a gene-of-interest. Dynamics in context also seem to be detectable, as well as gene-specific functions. *GeneCOCOA* offers a different approach to other methods, which appears to identify more biologically relevant gene functions than similar tools, although benchmarking these kinds of approaches remains highly challenging.

## Discussion

This manuscript describes *GeneCOCOA*, a method designed to implement both co-expression and functional enrichment analyses focused on a gene-of-interest (GOI). Evidence of the functionality of *GeneCOCOA* was demonstrated by using transcriptome profiling data arising from monogenic diseases, and identifying co-expressed gene sets with a relevant gene in each scenario. The use of *GeneCOCOA* to detect context-specific alterations in gene function was illustrated using RNA-sequencing data arising from a large cohort of patients suffering from acute myeloid leukemia. Here, functional gene sets associated with disease progression and prognosis could be found to be significantly co-expressed with *FLT3*, a known driver of the disease. The performance of *GeneCOCOA* relative to similar methods was compared across several distinct contexts, and showed that *GeneCOCOA* has the potential to fill a previously underpopulated niche in the toolkit of gene expression data analysis.

Advancements in next-generation sequencing technology have resulted in an abundance of high quality, publicly available transcriptome profiling data from a wide range of species, conditions and stimuli [3]. This has shifted the experimental bottleneck from data generation towards data analysis, with a resulting requirement for robust, efficient methods to extract maximal insight from these data. This must be accomplished whilst simultaneously maintaining ease-of-use for the user, many of whom are not expert computational biologists. Another by-product of this wealth of data is that researchers with specific genes-of-interest can query these data for metrics such as co-expression. However, manually curating co-expression results to derive biological insight can be complex and time-consuming.

Herein, we demonstrated that *GeneCOCOA*is capable of providing the user with functional gene sets which are enriched in their co-expression with a GOI. The functionality of *GeneCOCOA* in conjunction with data from large cohort experiments was demonstrated with a large data set consisting of 79 RNA-sequencing samples [36, 37], where the known functional role of *FLT3* could be recapitulated. In this illustrative example, the link between the gene-of-interest and experimental condition is extremely well established. This makes it difficult to truly assess the sensitivity of *GeneCOCOA* for discovering *de novo* functional roles of a GOI in a given condition.

In further illustrative use cases, *GeneCOCOA* was implemented on genes implicated as being causative for amyotrophic lateral sclerosis and familial hypercholesterolemia, specifically the GOIs *SOD1* [69] and *LDLR* [66]. In each case, *GeneCOCOA* identified functional, co-expressed enriched terms pertinent to the given disease. It should be noted, however, that in each case there were several replicates per condition (11 vs. 11, and 13 vs. 10, respectively). These replicate numbers are relatively uncommon in experimental setups designed around cell culture systems, where three biological replicates per biological condition is common [70]. The identification of robust enrichments when *GeneCOCOA* is provided with datasets of this smaller size is more challenging than when using larger datasets, and certainly represents a potential drawback of the approach. However, transcriptome profiling of larger patient cohorts is becoming increasingly common and accessible [71–73], providing ideal input for *GeneCOCOA* and similar tools. Another caveat to consider in the course of analysis of transcriptomic data with *GeneCOCOA* or any similar method, is the disconnect between expression and true function. Whilst *GeneCOCOA* is capable of using an array of curated gene annotation databases to infer potential functionality, a vast number of genes remain uncharacterized with regard to functional importance [74]. These genes are therefore excluded from the analysis, despite potentially interesting co-expression with the gene in question. Similarly, in a native co-expression analysis without any functional subsetting of genes, genes co-expressed with one another may in fact have diverse functions. For example, genes whose products make up negative feedback loops may be similarly regulated in order to provide a controlled response to a stimulus, despite having antagonistic functions [75]. In a systematic comparison of *GeneCOCOA* against similar methods (*GeneWalk*, *DAVID* and *Correlation AnalyzeR*), *GeneCOCOA* was able to identify a greater proportion of evidence-linked disease-relevant gene-GO term relationships. By computing these links across a number of diseases, it could be shown that disease-relevant associations reported by *GeneCOCOA* tended to be enriched in specificity for the diseases in question. However, it should be stated that making concrete conclusions on the relative performance of these types of methods is highly challenging, given the difficulties in ascribing true positive and true negative validation sets. This arises from the curated nature of gene sets, which rely wholly on published gene functions, as well as the extent and quality of databases used to record and document relationships between genes and functions. A consequence of this approach is that there may be genes not yet linked to a function or disease, which may just be unstudied in that capacity rather than irrelevant. For example, inflammatory genes such as *TNF* and *TGFB1* (both annotated as being important to myocardial infarction) are not included in the list of genes associated with Alzheimer’s disease on *DisGeNET*. As a consequence, significant associations reported for these genes (**Supp. Fig. S4**) with Alzheimer’s-relevant terms were marked as quasi-false positives. Yet, dysregulations related to these genes have been linked to the development of Alzheimer’s disease in prior research [76–79]. Similarly, *GeneCOCOA* also reported false positive associations in the amyotrophic lateral sclerosis (ALS) data set for the genes *BCL2* and *BAX*. While they are present in the Alzheimer’s disease gene set, these apoptotic genes have also been described as mediators of motor neuron loss in ALS [80–82]. Thus, the supposedly false positive associations returned by *GeneCOCOA* might, in several cases, hint at biologically meaningful GOI-disease associations which are not reflected in our strict approach to the definitions of ground truth.

From a methodological perspective, it was interesting that the relatively simple methods employed by *Correlation AnalyzeR* and *GeneCOCOA* both outperformed the more complex method implemented in *GeneWalk*. *Correlation Analyzer* ’s approach of considering entire gene sets in their enrichment analysis could result in a decreased sensitivity compared to *GeneCOCOA*, which samples subsets of gene sets. This would explain the greater sensitivity (but additionally increased false positive rate) of *GeneCOCOA*. The authors of *Correlation AnalyzeR* recommend input data with many samples in order for a robust analysis, whereas the iterative sampling approach of *GeneCOCOA* might permit increased performance on smaller datasets. What the performance of these two similar methods shows, is that using co-expression in combination with functional enrichment is a valid approach for inferring gene function, particularly of previously unstudied genes. Which specific method of co-expression analysis and functional enrichment should be used likely depends on the type and extent of the input data.

The formulation of *GeneCOCOA* to provide a functionally-resolved co-expression analysis framework is designed to minimize both data and time loss when moving data between different methods. Performance is largely determined by the iterative computation of background gene sets, the number of which may be set by the user. We aimed to maximize ease-of-use by formulating *GeneCOCOA* as an R [83] package, thereby making it simple to integrate the analysis with common workflows such as differential gene expression analysis [45, 84].

In summary, *GeneCOCOA* provides a method by which users can infer putative functions of a gene-of-interest based on co-expression of the given gene with curated sets of functionally-annotated genes. *GeneCOCOA* therefore empowers users to take advantage of the growing number of publicly available transcriptome profiling datasets, in order to provide greater functional insight and generate new hypotheses pertaining to the roles of individual genes in different contexts.

## Conclusion

- *GeneCOCOA* is a combined method for the identification of functional gene sets which are significantly co-expressed with a gene-of-interest.
- The method can be used in a highly flexible manner on user-supplied or publicly available transcriptome profiling data.
- Function gene sets can be provided by the user, or taken from curated, publicly available databases which hold information on ontologies, pathways and diseases.
- *GeneCOCOA* successfully recapitulates functional signatures of genes implicated in monogenic diseases.
- *GeneCOCOA* detects greater numbers of evidence-linked gene-disease relationships than similar methods.

## Authors contributions

S.Z., M.H.S. and T.W. conceived the idea for the study. R.P.B, M.S.L, S.W. and T.O. provided key data and input. S.Z. performed the analyses. S.Z. and T.W. wrote the code and formulated the manuscript. All listed authors read, commented on and edited the manuscript.

## Supporting information

**Figure S1.**
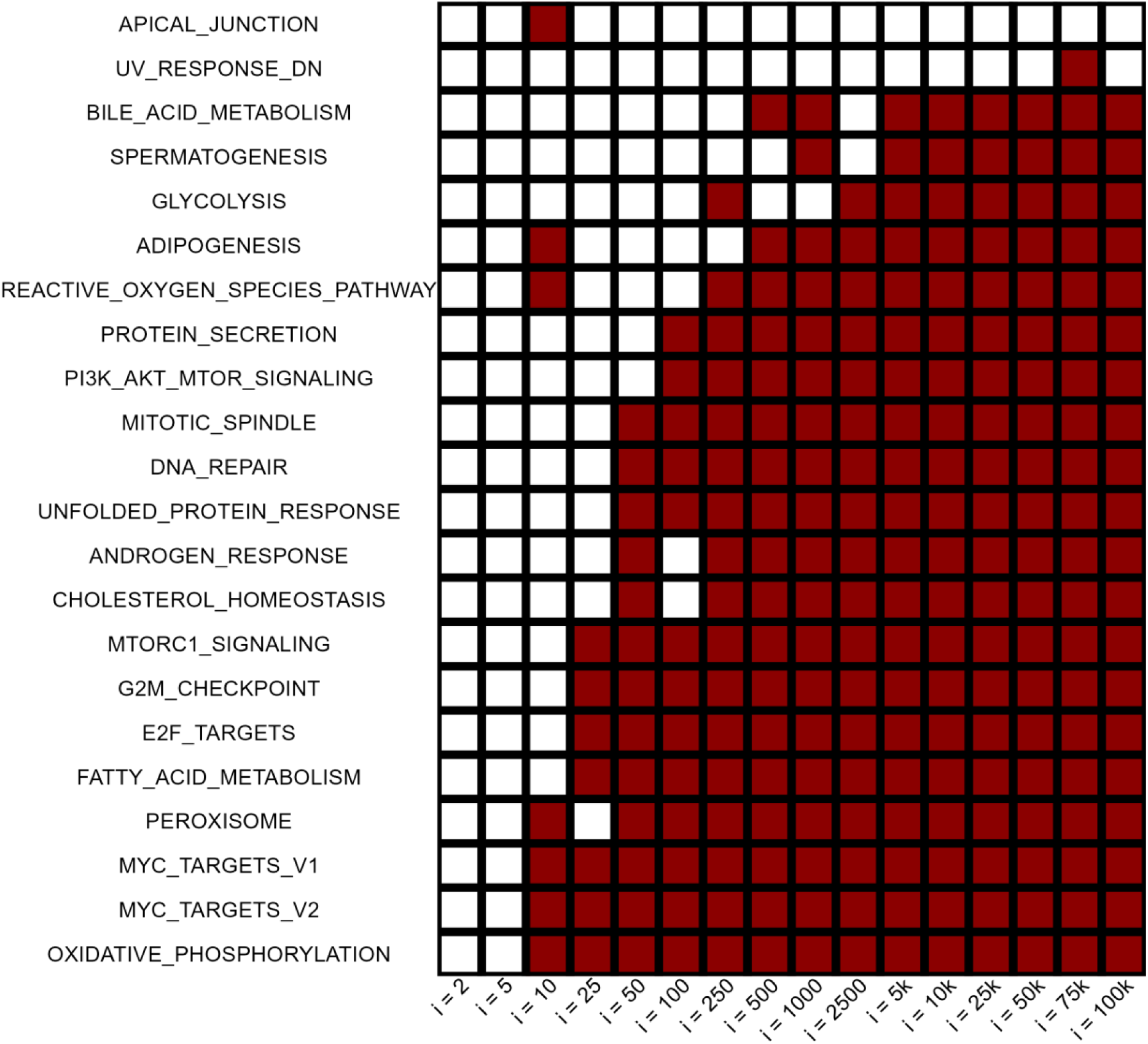
Identification of recommended number of bootstraps. With different values for number of bootstrapping rounds were tested, *i*=1000 was found to provide the best trade-off between efficiency and power. Displayed here are exemplary results for the association between *FLT3* and the 50 MSigDB hallmark gene sets in the expression data set of 136 AML patients. We inspected the results of 16 GeneCOCOA runs with bootstrap rounds ranging from 2 to 100k. All terms which were identified as significant (*P*_adj_<0.05) in any of the runs are listed as rows, while columns indicate the different GeneCOCOA runs. White tiles indicate that this term was not identified as significant in the respective GeneCOCOA run, while red indicates that it was returned as one of the terms significantly associated with *FLT3* expression.

**Figure S2.**
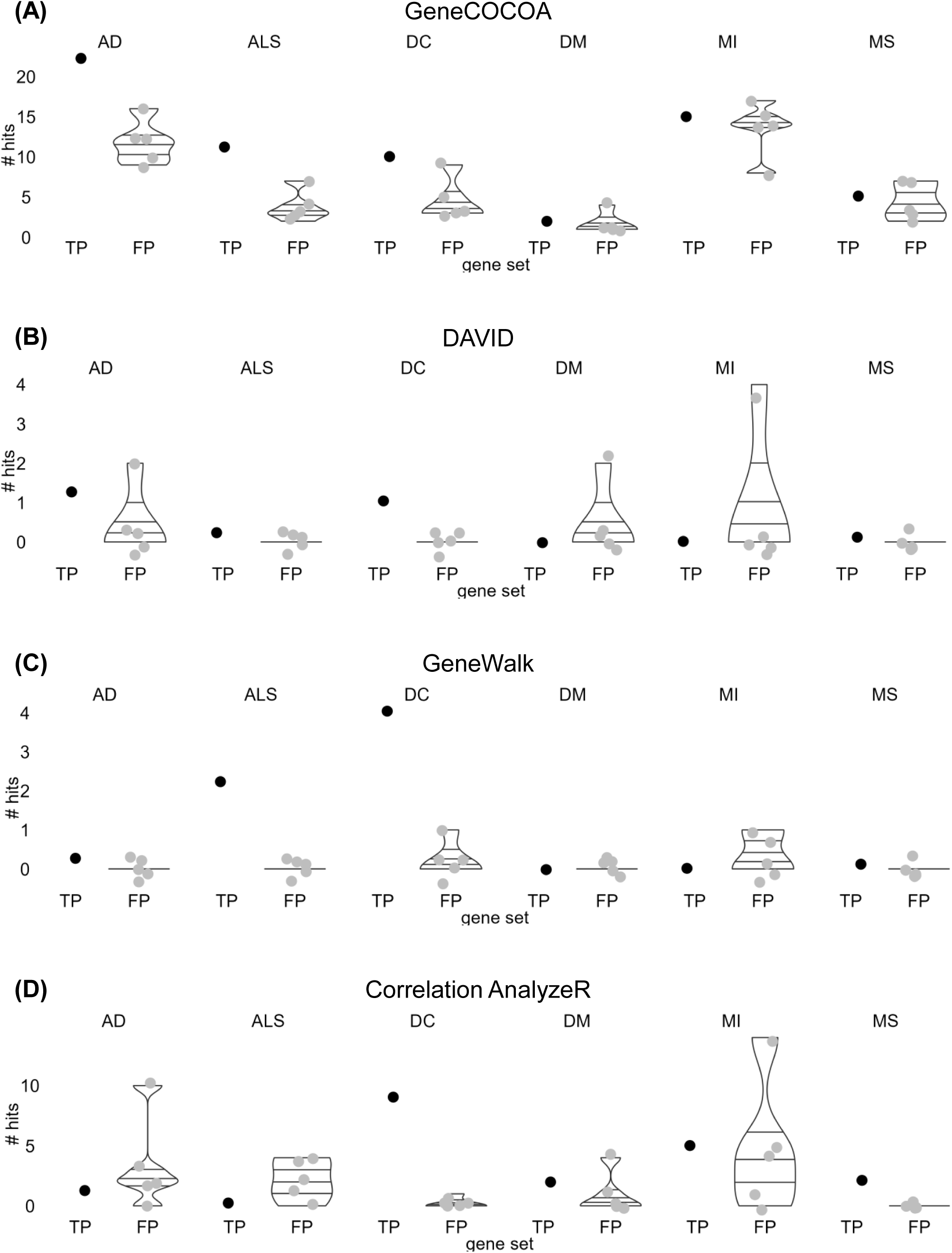
Comparison of true positives and false positives hits across gene sets. For each gene set, we evaluated the number of hits by method, differentiating true positives (TP; hits in the original disease context) from false positives (FP; hits in other disease contexts). **(A)** Across gene sets, the number of hits returned by GeneCOCOA in the TP condition is either higher or comparable to any other number of hits in FP contexts. DAVID and GeneWalk recover a smaller number of hits in general. While GeneWalk – except for the case of MI – manages to retain a good TP:FP ratio **(C),** DAVID **(B)** and Correlation AnalyzeR **(D)** report more FP than TP hits in a third of the cases.

**Figure S3.**
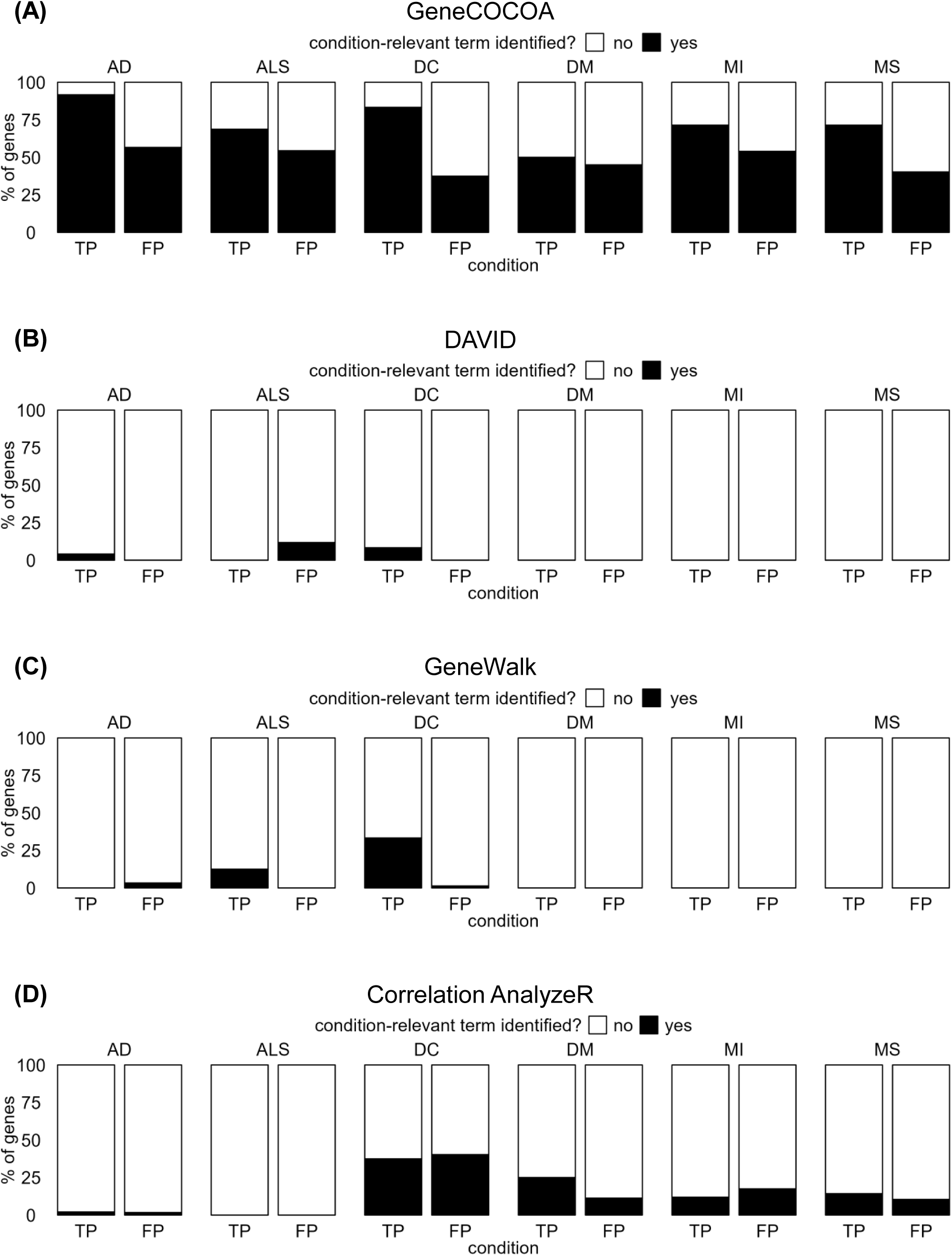
Comparison of true positives and false positives across conditions. For each condition, the set of genes which are disease-relevant as per DisGeNET can be defined as the *true* data set, all other genes are defined as *other*. **(A)** Comparing the proportions of *true* genes with disease-relevant term hits against the proportion of *other* genes with disease-relevant term hits, GeneCOCOA consistently manages to recover more *true* hits than *other* hits across all conditions. **(B)** DAVID and **(C)** GeneWalk show only a negligible proportion of *other* hits. Yet, these methods also fail to recover a substantial amount of *true* hits. (D) In two cases, Correlation AnalyzeR shows slightly more *true* than *other* hits. Yet, in all other cases there are at least as many *other* as *true* hits. The overall percentage of *true* hits recovered is smaller than in the GeneCOCOA runs.

**Figure S4.**
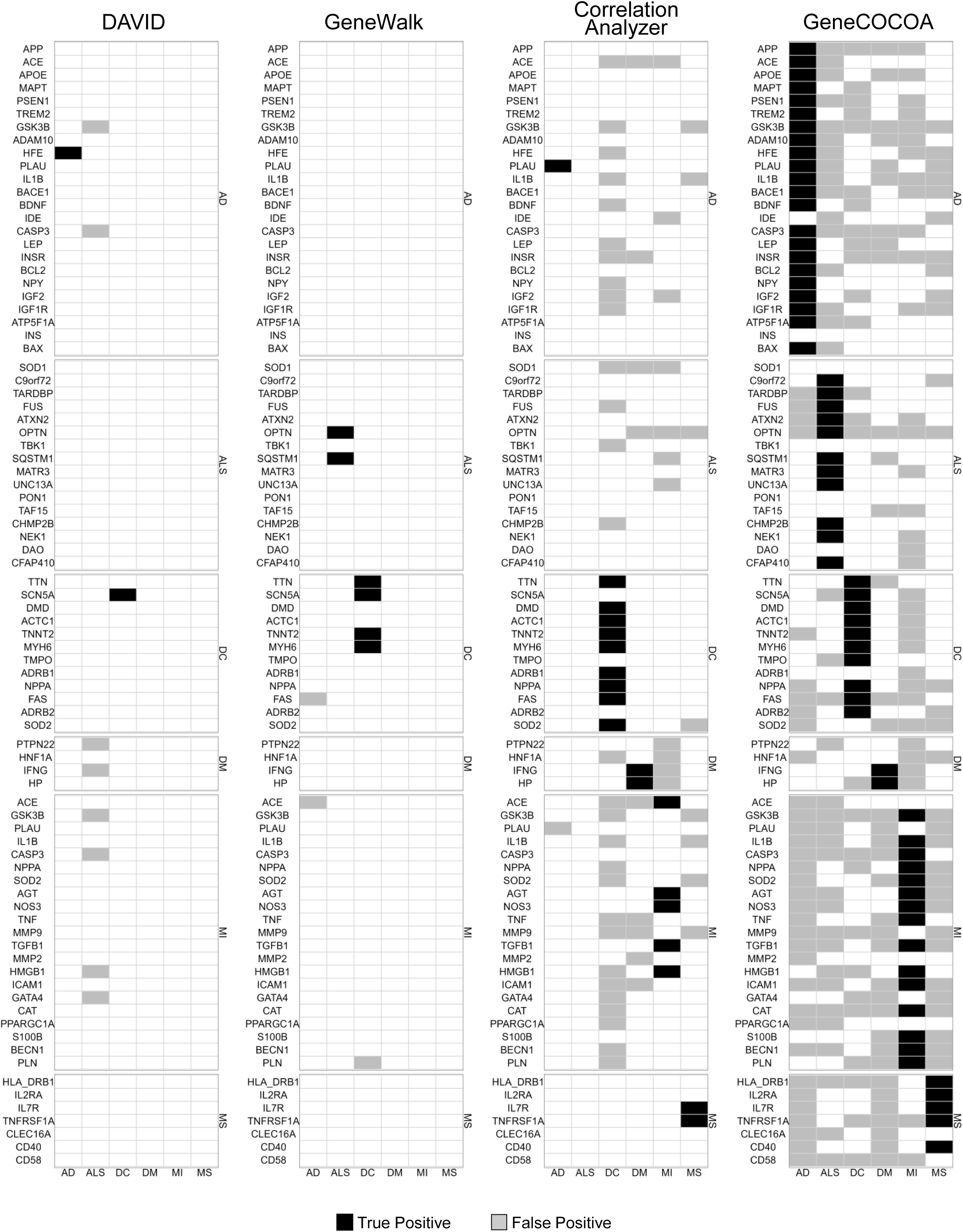
False/true positive matrices for all three methods with gene symbols. Summary of true positive and false positive gene-term associations per set of disease-relevant genes across all diseases, as computed by DAVID, GeneWalk, Correlation AnalyzeR and GeneCOCOA.

